# A benchmarking of human Y-chromosomal haplogroup classifiers from whole-genome and whole-exome sequence data

**DOI:** 10.1101/2022.09.19.508481

**Authors:** Víctor García-Olivares, Adrián Muñoz-Barrera, Luis A. Rubio-Rodríguez, David Jáspez, Ana Díaz-de Usera, Antonio Iñigo Campos, Krishna R. Veeramah, Santos Alonso, Mark G. Thomas, José M. Lorenzo-Salazar, Rafaela González-Montelongo, Carlos Flores

## Abstract

The non-recombinant region of the Y chromosome (NRY) contains a great number of polymorphic markers that allows to accurately reconstruct pedigree relationships and retrieve ancestral information from study samples. The analysis of NRY is typically implemented in anthropological, medical, and forensic studies. High-throughput sequencing (HTS) has profoundly increased the identification of genetic markers in the NRY genealogy and has prompted the development of automated NRY haplogroup classification tools. Here, we present a benchmarking study of five command-line tools for NRY haplogroup classification. The evaluation was done using empirical short-read HTS data from 50 unrelated donors using paired data from whole-genome sequencing (WGS) and whole-exome sequencing (WES) experiments. Besides, we evaluate the performance of the top-ranked tool in the classification of data of third generation HTS obtained from a subset of donors. Our findings demonstrate that WES can be an efficient approach to infer the NRY haplogroup, albeit generally providing a lower level of genealogical resolution than that recovered by WGS. Among the tools evaluated, YLeaf offers the best performance for both WGS and WES applications. Finally, we demonstrate that YLeaf is able to correctly classify all samples sequenced with nanopore technology from long noisy reads.

## 1. Introduction

The Y chromosome (chrY) is one of the smallest chromosomes in the human genome (~60 Mb). A high proportion of this chromosome (95%), known as the non-recombining region (NRY), shows a patrilineal inheritance following a haploid behavior due to its lack of chromosomal recombination during meiosis (Quintana-Murci & Fellous, 2001). Because of this, the NRY allows for a precise reconstruction of its genealogy back to a common ancestor, as described by the coalescence theory (Underhill & Kivisild, 2007). The study of NRY has a wide range of applications in fields such as evolutionary anthropology and population history (Pinotti et al., 2019; Zeng et al., 2018), medical genetics (Colaco & Modi, 2018; Grassmann et al., 2019), and forensic science (Kayser, 2017; Zhou et al., 2022).

The advent of high-throughput sequencing (HTS) technology has brought about a revolution in the development of human genomics and medicine. The decrease in costs and the increase in coverage of both whole-genome sequencing (WGS) and whole-exome sequencing (WES) applications offer the possibility of improving the chrY research through deeper and more optimal analyses (Anderson et al., 2019; Levy & Myers, 2016). However, this chromosome presents regions that are challenging to sequence, such as short tandem repeats (STRs) (Alvarez-Cubero et al., 2018; Charlesworth, 2003). In this regard, third-generation HTS with the development of longer reads with greater read depth may help to improve the mapping of such complex repeat sequences (Anderson et al., 2019; Jobling & Tyler-Smith, 2017). An example of this is the use of nanopore technology (ONT, Oxford Nanopore Technologies, Oxford, UK) to successfully generate the first African human chrY reference assembly (Kuderna et al., 2019).

The use of HTS technology for human genome sequencing has enabled the discovery of new variants in the chrY providing a remarkable increase in the volume of marker information available to trace the human paternal lineages (Claerhout et al., 2021). In this regard, the International Society of Genetic Genealogy (ISOGG-Y-DNA tree; https://isogg.org/tree/) has compiled since 2006 all new variants in the NRY, generating a database that currently hosts more than 90,650 unique biallelic variants. The NRY diversity has been structured following a phylogenetic hierarchy based on variants that define distinct clades representing haplotypes commonly referred to as haplogroups (The Y Chromosome Consortium, 2002). The study of these haplogroups allows to trace origins, patterns of differentiation between populations, and to unravel historical patterns of human migration over time (Calafell & Larmuseau, 2017). Haplogroup identification is, therefore, a key step in recovering pedigree relationships and ancestral information from analyzed samples.

The exponential increase in the number of NRY markers, which concomitantly associates with a rise in the complexity of the chrY tree, and the development of HTS imposes a bioinformatics challenge for inferring the patrilineal genetic genealogy of the study samples. To take advantage of the potential offered by HTS technology, the number of automated NRY classification tools has seen a considerable increase in recent years (Chen et al., 2021; David Poznik, 2016; Jagadeesan et al., 2020; Martiniano et al., 2022; Ralf et al., 2018, 2019; Van Geystelen et al., 2013; Zhang et al., 2013). However, an unmet need to date is a comparative study evaluating the performance of each tool. Here we present a benchmarking analysis of several command-line tools for automated human NRY classification using empirical short-read HTS data from two of the most widely used applications in human genetics, WGS and WES. In addition, we assessed the performance of the best performing tool in third-generation HTS long noisy read WGS data obtained by nanopore technology for a subset of the donors.

## 2. Materials and Methods

### Samples, library preparation, and sequencing

The study was approved by the Research Ethics Committee of the Hospital Universitario Nuestra Señora de Candelaria (CHUNSC_2020_95) and performed according to The Code of Ethics of the World Medical Association (Declaration of Helsinki).

Fifty DNA samples from unrelated donors were used for the study after informed consent. All samples were sequenced in parallel using short-read WGS and WES. The construction of libraries was performed with Illumina preparation kits (**Table S1**) following the manufacturer’s recommendations (Illumina Inc., San Diego, CA, USA). The Nextera DNA Prep kit and Illumina DNA Prep were used for WGS. The same samples were processed with Nextera DNA Exome and Illumina DNA Prep with Enrichment as described elsewhere (Díaz-de Usera et al., 2020). The library quality controls were carried out in a TapeStation 4200 (Agilent Technologies, Santa Clara, CA, USA) and sequencing was conducted on HiSeq 4000 or NovaSeq 6000 (Illumina, Inc., San Diego, CA, USA) instruments.

Seven of these samples were also sequenced using long noisy read WGS data obtained with nanopore technology at KeyGene (Wageningen, The Netherlands). Sequencing was performed on a PromethION system (ONT) for 64 h using one FLO_PR002 (R9.4.1 pore) flow cell per sample following the manufacturer’s recommendations. Basecalling was conducted on the PromethION computing module using MinKNOW v1.14.2 with Guppy v2.2.2, and data preprocessing metrics were carried out with PycoQC v2.5.2 (Leger & Leonardi, 2019).

### Bioinformatic processing

Processing of short-read WGS and WES data was carried out using an in-house pipeline based on GATK v4.1 for WGS and GATK v3.8 for WES (McKenna et al., 2010). Raw reads were assessed using FastQC v.0.11.8 software (Andrews, 2010) and aligned to the GRCh37/hg19 reference genome using BWA-MEM v0.7.15 (Li & Durbin, 2009). Quality control of aligned reads was performed with Qualimap v2.2.1 (García-Alcalde et al., 2012). The alignments were then processed for duplicate marking and base quality score recalibration (DePristo et al., 2011). Variant calling step was conducted by GATK HaplotypeCaller following the Broad Institute’s best practices workflow for germline short variant discovery. From the resulting BAM files, the NRY region (2.64-59.03 Mb) was extracted by using SAMtools v1.12 (Li et al., 2009). Regarding the WES data, it was observed that of the 676 DNA capture probes used for hybridisation-based target enrichment of the chrY regions, 90% are within the NRY, covering 0.22% of this region. For ONT data, raw long noisy reads were first preprocessed with FiltLong v0.2.1 (https://github.com/rrwick/Filtlong) to filter reads shorter than 1,000 bp. Filtered reads were assessed using NanoPlot v1.38.1 (De Coster et al., 2018), then aligned to the GRCh37/hg19 reference genome using Minimap2 v2.22-r1101 (Li, 2018) and sorted with SAMtools v1.14, extracting only reads aligned to the NRY region. All these bioinformatic processes were computed at TeideHPC infrastructure (https://teidehpc.iter.es/en/home/).

### Sex quality control

To identify the sex of the donnors, a quality control was performed based on both the self-reported sex of the individual and two bioinformatics approaches. The first approach, performed by Somalier v0.2.15 (Pedersen et al., 2019), identifies the sex of the sample from the depth of the X and Y chromosome reads. For the second approach, an in-house heuristic script (https://github.com/genomicsITER/sexQC-for-NGS-data) was used. The analysis involves assessing the depth of 11 genes in the non-pseudoautosomal regions of the X and Y chromosomes using high quality mapped reads (MappingQuality>50). All samples used for this study were identified as males by consensus from these two approaches.

### Y-chromosomal haplogroup classification

Among the tools available from the literature, we selected eight tools that were open-source and offered a command-line interface (**Table 1**). These were run with the 2020 version (v15.73) of the ISOGG repository database, which contains more than 90,000 polymorphic markers and it constitutes the central reference for many bioinformatic tools to classify human chrY sequences. However, three of the tools (YHap, AMY-tree, and yhaplo) were finally excluded from the study because they imposed limitations for database updates. For YLeaf, the version 2.2 was used since the newer 3.1 version does not use the ISOGG marker identifiers in the classification. Y-Lineage Tracker has the option of using VCF and BAM input files, fostering an evaluation with the two alternative supported files. The haplogroup classification process was executed using a workstation running CentOS 7 with 2 Intel Xeon Cascade Lake 6252 Gold CPUs at 2.1 GHz and 384 GB of RAM. All tools were run using the default parameters.

**Table 1.**
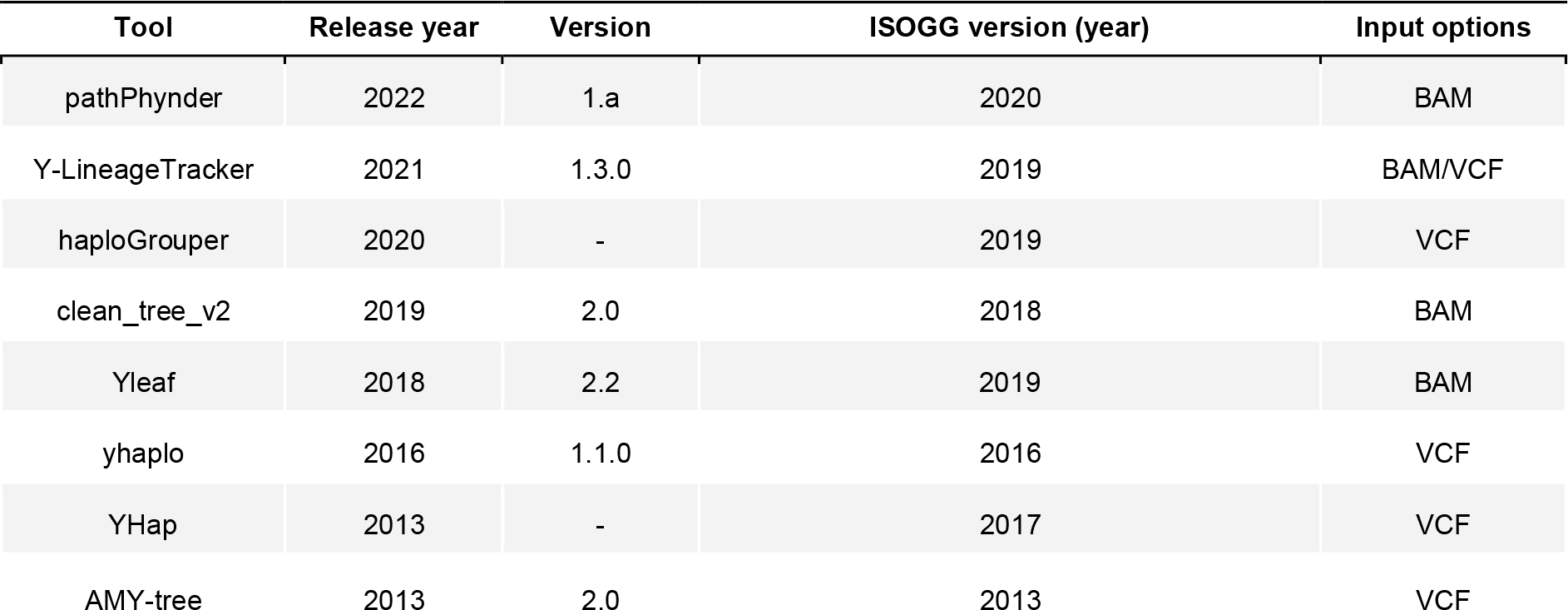
List of tools assessed for human Y-chromosomal haplogroup classification. All the tools assessed classify according to the ISOGG nomenclature by using the latest version 15.73 (2020).

| Tool | Release year | Version | ISOGG version (year) | Input options |
| --- | --- | --- | --- | --- |
| pathPhynder | 2022 | 1.a | 2020 | BAM |
| Y-LineageTracker | 2021 | 1.3.0 | 2019 | BAM/VCF |
| haploGrouper | 2020 | - | 2019 | VCF |
| clean_tree_v2 | 2019 | 2.0 | 2018 | BAM |
| Yleaf | 2018 | 2.2 | 2019 | BAM |
| yhaplo | 2016 | 1.1.0 | 2016 | VCF |
| YHap | 2013 | - | 2017 | VCF |
| AMY-tree | 2013 | 2.0 | 2013 | VCF |

Unlike WGS, which recovers a larger part of the NRY, WES only partially recovers the NRY (**Table S1**). This difference may lead to discrepancies in the haplogroup classification obtained by both applications simply because it is expected that a lower level of resolution could be obtained for WES in any given sample. To deal with this limitation in the benchmarking, we took as the reference for the comparisons the maximum classification level retrieved by WES that matches the one obtained from the WGS data.

## 3. Results

### Sequencing summary for short-read and long-read sequencing

The mean (± SD) number of NRY reads recovered per sample (n = 50) for short-read WGS and WES data were 8,329,867 ± 2,460,724 and 575,090 ± 176,201, respectively (**Table S1**). For WGS, 33.66% of the NRY was covered at least at 10X. For WES, this percentage decreased to 0.9%. However, if only the exonic regions of NRY are taken into account, 84.46% was covered by WES at least at 10X. The mean (± SD) depth of coverage recovered across the NRY region for WGS was 13X ± 4 (range: 6-28X). The depth decreased to less than 1X for WES, although it was as high as 67X ± 19 (range: 27-111X) when only the exonic regions were included in the analysis. As for the detected single nucleotide variants (SNVs), the mean (± SD) depth of coverage per SNV call had a value of 60 ± 17 in WGS, decreasing to 32 ± 15 in WES. For ONT, the mean (± SD) number of NRY reads per sample (n = 7) recovered was 177,436 ± 40,374. The mean NRY depth of coverage was 11X ± 2 (range: 8-14X) and 31.57% of the NRY was covered at least at 10X (**Table S1**). Furthermore, while the WGS data from both sequencing technologies provided a homogeneous depth of coverage profile across the NRY region (except in regions adjacent to the centromere and the heterochromatic region because of their complexity), the WES data showed a heterogeneous profile with enriched sites (peaks) associated to the capture of exons that are embedded within undetected regions (**Figure 1**).

**Figure 1.**
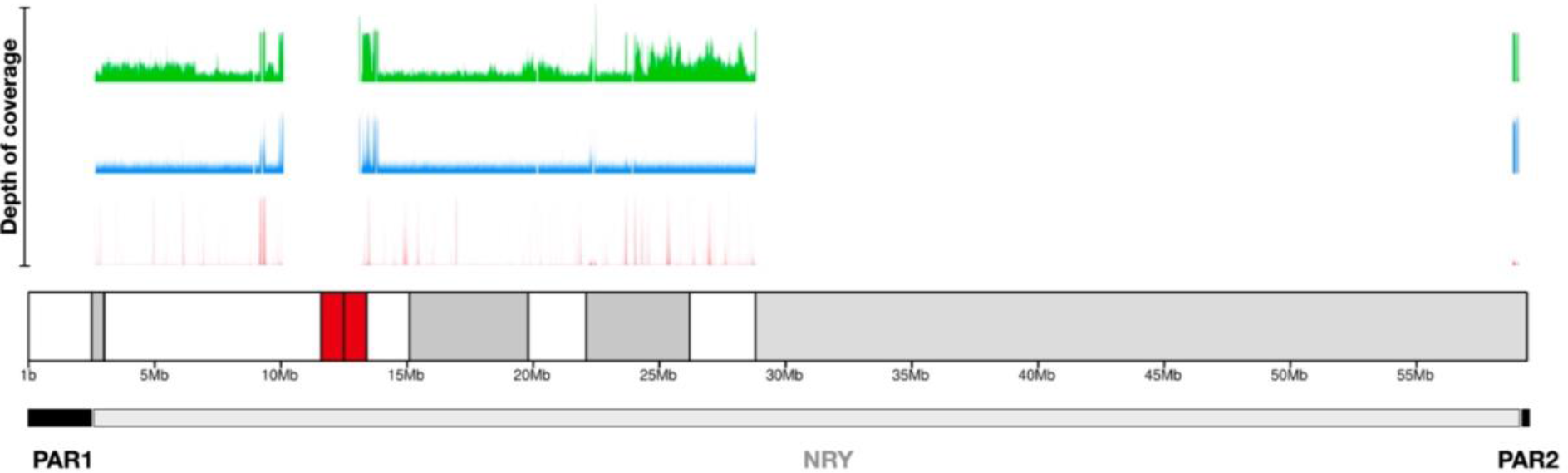
Plot of the depth of coverage for short-read and long-read sequencing in the non-recombining portion of the Y chromosome (NRY) of an exemplar sample. Long-read WGS data is shown in green. Short-read WGS and WES data are colored in blue and red, respectively. In the ideogram of the Y chromosome, the heterochromatic regions (positive C-band) and the centromere are colored in gray and red, respectively. In the lowest panel, the pseudoautosomal regions (PAR1 and PAR2) and the NRY are represented in black and gray, respectively. To harmonize the results obtained from the three approaches, the depth of coverage was normalized to 100X. The R package karyotypeR v1.2.2 (Gel & Serra, 2017) was used to generate the depth of coverage plot.

### Consensus haplogroup classification

Based on the metrics retrieved for the short-read data, WGS showed higher values than WES for both the breadth and depth of coverage parameters, both of which are closely related to higher statistical support for variant detection. Therefore, we establish the WGS-derived haplogroup as the ground truth. To assign the haplogroup of each sample, the haplogroup most frequently classified by all the tools assessed was used as consensus. Fourteen samples showed 100% concordance among the tools evaluated, and the remaining, we obtained a mean concordance rate of 66.20% (**Table S2**). However, in most cases where discordance was reported, they were due to distinct haplogroup level classifications and not to misclassification. In four of the samples inconsistencies among the tools precluded a straightforward indication of the consensus haplogroup. In these four cases, the classification result of YLeaf was used as the ground truth given its higher classification accuracy demonstrated in all other samples. Considering only the WGS results, YLeaf offered the highest classification accuracy (94%). With slightly lower, but relatively high performance (>70%), we found clean_tree_v2, LineageTracker (with VCF as the input file), Haplogrouper, and pathPhynder. The worst performing tool was LineageTracker using BAM as the input file since it misclassified 56% of the samples. Based on the limitations outlined in the methodology and in order to harmonize the results, the consensus haplogroup per sample used for the benchmarking was subordinated to the maximum level of resolution retrieved by WES (**Table S2**).

### Haplogroup classification

The classification accuracy provided by short-read WGS data reached an average of 88.67%, while for WES it decreased to an average of 43.67% (**Table S2**). On average (± SD), there were less discordances on the WGS classifications (0.68 ± 0.96) than for WES (3.38 ± 1.01). By the classification accuracy for WGS data, LineageTracker (VCF as input file) and YLeaf showed the highest accuracy, classifying precisely 98% and 96% of the analyzed samples, respectively. With slightly lower accuracy were the following tools: pathPhynder (92%), Haplogrouper (90%), and clean_tree_v2 (88%). LineageTracker, using BAM as input file, was the least accurate tool for WGS data, misclassifying 32% of the samples. For WES data, YLeaf showed the highest classification accuracy among all tools, classifying precisely 88% of the analyzed samples. With a slightly lower accuracy than YLeaf were clean_tree_v2 and pathPhynder, with an average of 80%. Haplogrouper and LineageTracker (VCF as input file) were the least accurate tools, yielding an incorrect haplogroup classification in more than 88% of the samples. LineageTracker from a BAM file did not provide a result for any of the samples. Since YLeaf showed the best performance for both WGS and WES short-read data, this tool was used to classify samples from ONT data. The seven samples sequenced with long noisy ONT reads were correctly classified at the haplogroup level retrieving the same classification resolution as the WGS short-read data.

## 4. Conclusions

The advent of HTS technologies, the notable increase in the number of NRY polymorphic markers detected, together with the importance of recovering ancestral information and pedigree relationships of study samples, have motivated the development of new automated classification tools to adapt to these challenges. In this study, we present a benchmarking of five classification tools that could be easily upgraded to new versions of the ISOGG-Y-DNA tree. The comparison was carried out with empirical paired HTS data from WGS and WES, two of the most widely used applications in human genetics and medicine. Our results support that WES provides sufficient information to classify the NRY haplogroup. Besides, we demonstrate that YLeaf shows the best performance among all the tools evaluated for both applications, although with a slight loss in classification accuracy on the WES data. In most of the samples, the classification retrieved matched with that inferred by WGS, although in several samples a relatively lower level of accuracy was observed. However, considering that WES sequences include a limited fraction of the NRY region, the performance achieved by YLeaf tool for WES data is remarkable. Regarding third-generation HTS, our findings show that, despite the lower *per-base* accuracy currently offered by the assessed technology, it did not preclude an equally accurate classification as that obtained from short-read data.

## 5. Acknowledgements

We would like to thank the support from our colleagues from the Teide-HPC Supercomputing facility (http://teidehpc.iter.es/en), which was funded by INP-2011-0063-PCT-430000-ACT (INNPLANTA program) from the Spanish Ministry of Economy and Competitiveness.

## 6. Author Contributions

Conceptualization, V.G.O., A.M.B., L.A.R.R., C.F.; Data curation, V.G.O., A.M.B., L.A.R.R., A.M.B., J.M.L.S., D.J.T., R.G.M. K.R.V., M.G.T.; Methodology, V.G.O., A.M.B., J.M.L.S., L.A.R.R., A.D.U., D.J.T., A.I.C., R.G.M.; Supervision, C.F.; Writing—original draft preparation, V.G.O., A.M.B., C.F.; Writing—review and editing, V.G.O., A.M.B., L.A.R.R., S.A., C.F. All authors have read and agreed to the published version of the manuscript.

## Supplementary Materials

**Table S1.**
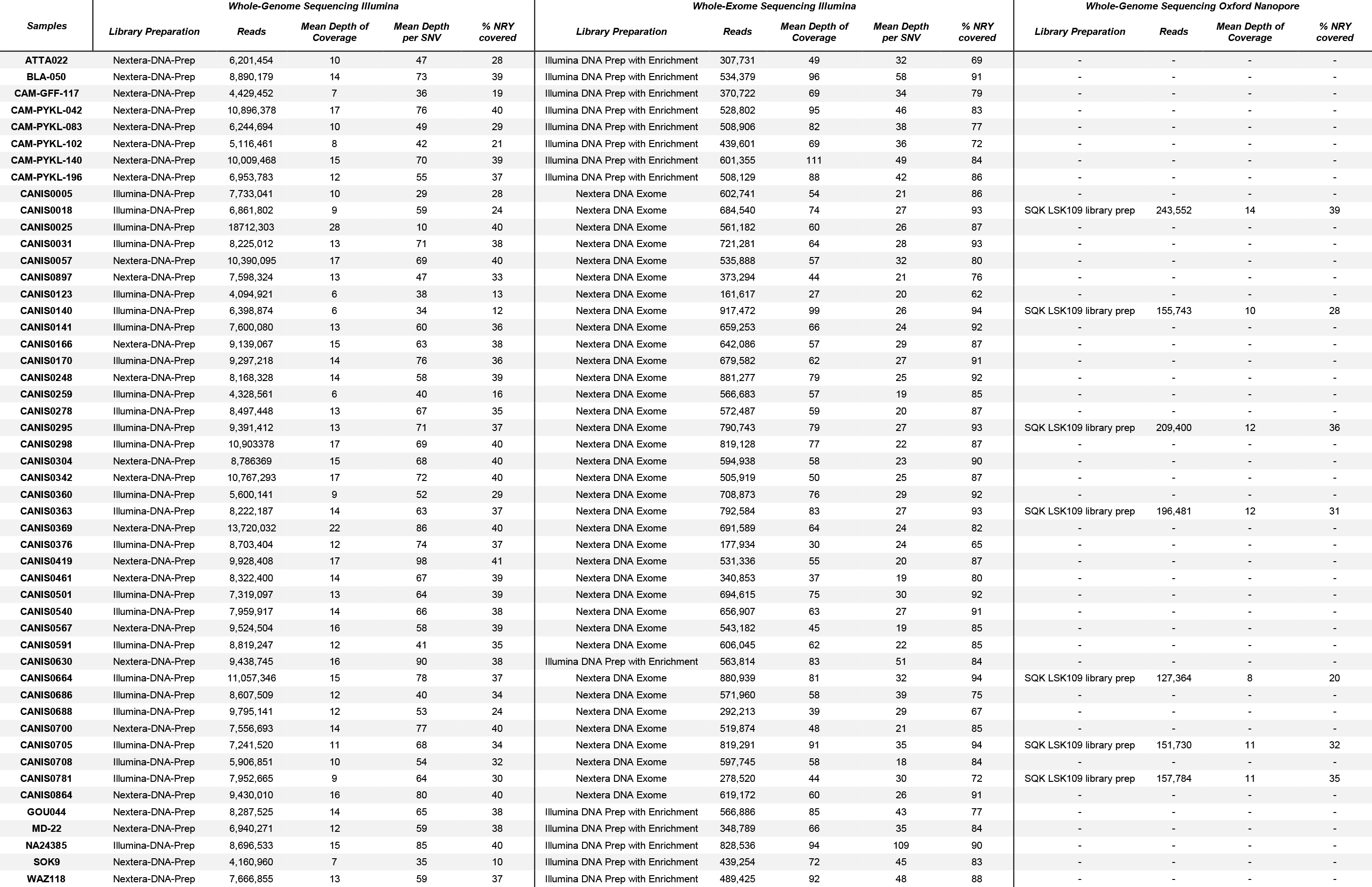
Short-read and long-read sequencing summaries of the samples.

**Table S2.**
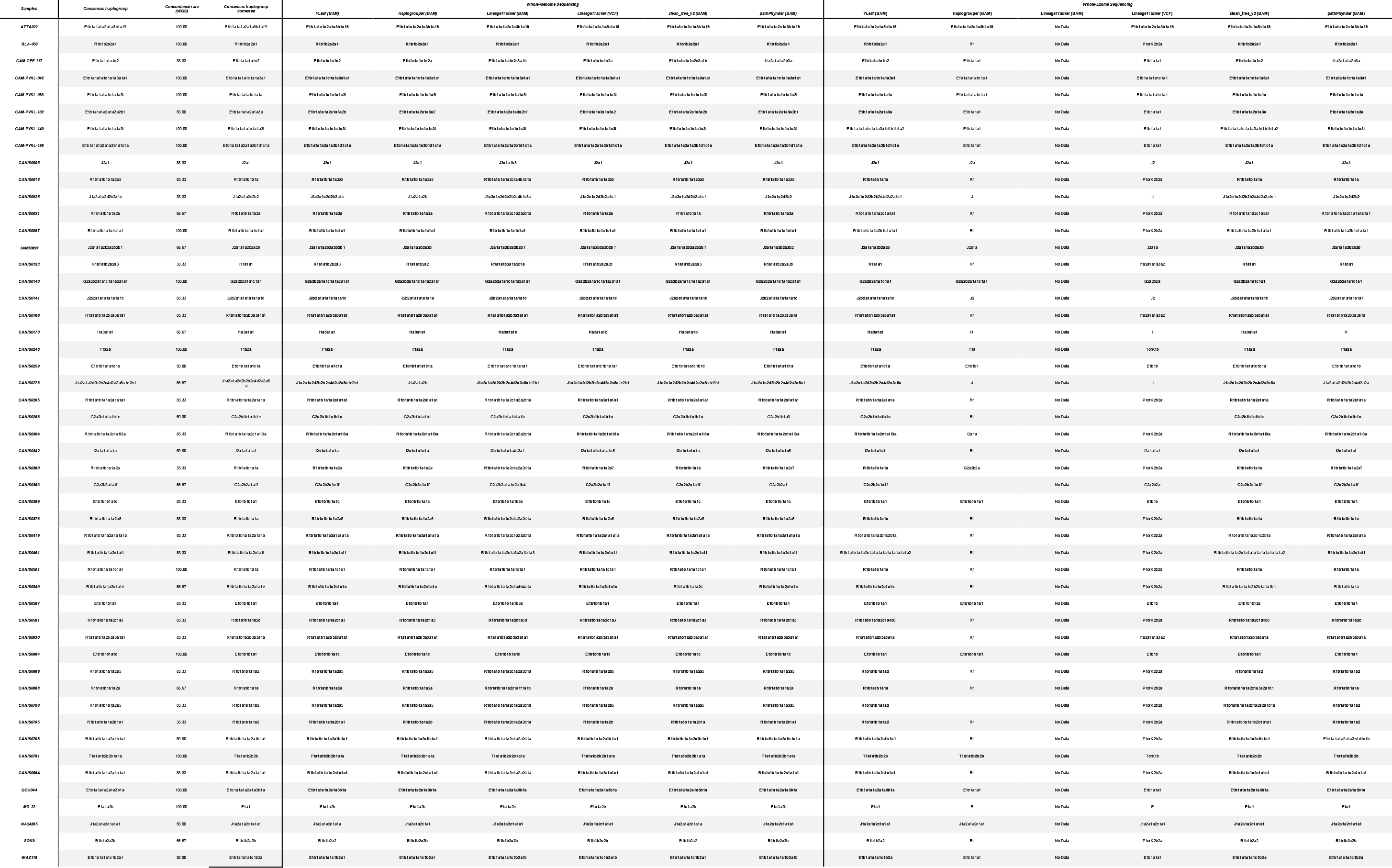
Per sample Y chromosome haplogroup classifications for the five tools based on short-read data. The retrieved classification that matches the consensus is shown in bold. A) WGS and B) WES datasets.

## References

Alvarez-Cubero, M. J., Santiago, O., Martínez-Labarga, C., Martínez-García, B., Marrero-Díaz, R., Rubio-Roldan, A., Pérez-Gutiérrez, A. M., Carmona-Saez, P., Lorente, J. A., & Martinez-Gonzalez, L. J. (2018). Methodology for Y Chromosome Capture: A complete genome sequence of Y chromosome using flow cytometry, laser microdissection and magnetic streptavidin-beads. Scientific Reports, 8(1), 9436.

Anderson, K., Cañadas-Garre, M., Chambers, R., Maxwell, A. P., & McKnight, A. J. (2019). The Challenges of Chromosome Y Analysis and the Implications for Chronic Kidney Disease. Frontiers in Genetics, 10, 781.

Andrews, S. (2010). FastQC: a quality control tool for high throughput sequence data. Babraham Bioinformatics, Babraham Institute, Cambridge, United Kingdom.

Calafell, F., & Larmuseau, M. H. D. (2017). The Y chromosome as the most popular marker in genetic genealogy benefits interdisciplinary research. Human Genetics, 136(5), 559–573.

Charlesworth, B. (2003). The organization and evolution of the human Y chromosome. Genome Biology, 4(9), 226.

Chen, H., Lu, Y., Lu, D., & Xu, S. (2021). Y-LineageTracker: a high-throughput analysis framework for Y-chromosomal next-generation sequencing data. In BMC Bioinformatics (Vol. 22, Issue 1). https://doi.org/10.1186/s12859-021-04057-z

Claerhout, S., Verstraete, P., Warnez, L., Vanpaemel, S., Larmuseau, M., & Decorte, R. (2021). CSYseq: The first Y-chromosome sequencing tool typing a large number of Y-SNPs and Y-STRs to unravel worldwide human population genetics. PLoS Genetics, 17(9), e1009758.

Colaco, S., & Modi, D. (2018). Genetics of the human Y chromosome and its association with male infertility. Reproductive Biology and Endocrinology: RB&E, 16(1), 14.

David Poznik, G. (2016). Identifying Y-chromosome haplogroups in arbitrarily large samples of sequenced or genotyped men. In Cold Spring Harbor Laboratory (p. 088716). https://doi.org/10.1101/088716

De Coster, W., D’Hert, S., Schultz, D. T., Cruts, M., & Van Broeckhoven, C. (2018). NanoPack: visualizing and processing long-read sequencing data. Bioinformatics, 34(15), 2666–2669.

DePristo, M. A., Banks, E., Poplin, R., Garimella, K. V., Maguire, J. R., Hartl, C., Philippakis, A. A., del Angel, G., Rivas, M. A., Hanna, M., McKenna, A., Fennell, T. J., Kernytsky, A. M., Sivachenko, A. Y., Cibulskis, K., Gabriel, S. B., Altshuler, D., & Daly, M. J. (2011). A framework for variation discovery and genotyping using next-generation DNA sequencing data. Nature Genetics, 43(5), 491–498.

García-Alcalde, F., Okonechnikov, K., Carbonell, J., Cruz, L. M., Götz, S., Tarazona, S., Dopazo, J., Meyer, T. F., & Conesa, A. (2012). Qualimap: evaluating next-generation sequencing alignment data. Bioinformatics, 28(20), 2678–2679.

Gel, B., & Serra, E. (2017). karyoploteR: an R/Bioconductor package to plot customizable genomes displaying arbitrary data. In Bioinformatics (Vol. 33, Issue 19, pp. 3088–3090). https://doi.org/10.1093/bioinformatics/btx346

Grassmann, F., Kiel, C., den Hollander, A. I., Weeks, D. E., Lotery, A., Cipriani, V., Weber, B. H. F., & International Age-related Macular Degeneration Genomics Consortium (IAMDGC). (2019). Y chromosome mosaicism is associated with age-related macular degeneration. European Journal of Human Genetics: EJHG, 27(1), 36–41.

Jagadeesan, A., Ebenesersdóttir, S. S., Guðmundsdóttir, V. B., Thordardottir, E. L., Moore, K. H. S., & Helgason, A. (2020). HaploGrouper: A generalized approach to haplogroup classification. Bioinformatics. https://doi.org/10.1093/bioinformatics/btaa729

Jobling, M. A., & Tyler-Smith, C. (2017). Human Y-chromosome variation in the genome-sequencing era. Nature Reviews. Genetics, 18(8), 485–497.

Kayser, M. (2017). Forensic use of Y-chromosome DNA: a general overview. Human Genetics, 136(5), 621–635.

Kuderna, L. F. K., Lizano, E., Julià, E., Gomez-Garrido, J., Serres-Armero, A., Kuhlwilm, M., Alandes, R. A., Alvarez-Estape, M., Alioto, T., Gut, M., Gut, I., Schierup, M. H., Fornas, O., & Marques-Bonet, T. (2019). Selective single molecule sequencing and assembly of a human Y chromosome of African origin. Nature Communications. https://doi.org/10.1101/342667

Leger, A., & Leonardi, T. (2019). pycoQC, interactive quality control for Oxford Nanopore Sequencing. Journal of Open Source Software, 4(34), 1236.

Levy, S. E., & Myers, R. M. (2016). Advancements in Next-Generation Sequencing. Annual Review of Genomics and Human Genetics, 17, 95–115.

Li, H. (2018). Minimap2: pairwise alignment for nucleotide sequences. Bioinformatics, 34(18), 3094–3100.

Li, H., & Durbin, R. (2009). Fast and accurate short read alignment with Burrows–Wheeler transform. Bioinformatics, 25(14), 1754–1760.

Li, H., Handsaker, B., Wysoker, A., Fennell, T., Ruan, J., Homer, N., Marth, G., Abecasis, G., Durbin, R., & 1000 Genome Project Data Processing Subgroup. (2009). The Sequence Alignment/Map format and SAMtools. Bioinformatics, 25(16), 2078–2079.

Martiniano, R., De Sanctis, B., Hallast, P., & Durbin, R. (2022). Placing Ancient DNA Sequences into Reference Phylogenies. Molecular Biology and Evolution, 39(2). https://doi.org/10.1093/molbev/msac017

Pedersen, B. S., Bhetariya, P. J., Brown, J., Marth, G., Jensen, R. L., Bronner, M. P., Underhill, H. R., & Quinlan, A. R. (2019). Somalier: rapid relatedness estimation for cancer and germline studies using efficient genome sketches. In bioRxiv (p. 839944). https://doi.org/10.1101/839944

Pinotti, T., Bergström, A., Geppert, M., Bawn, M., Ohasi, D., Shi, W., Lacerda, D. R., Solli, A., Norstedt, J., Reed, K., Dawtry, K., González-Andrade, F., Paz-Y-Miño, C., Revollo, S., Cuellar, C., Jota, M. S., Santos, J. E., Jr, Ayub, Q., Kivisild, T., … Tyler-Smith, C. (2019). Y Chromosome Sequences Reveal a Short Beringian Standstill, Rapid Expansion, and early Population structure of Native American Founders. Current Biology: CB, 29(1), 149–157.e3.

Quintana-Murci, L., & Fellous, M. (2001). The Human Y Chromosome: The Biological Role of a “Functional Wasteland.” Journal of Biomedicine & Biotechnology, 1(1), 18–24.

Ralf, A., Montiel González, D., Zhong, K., & Kayser, M. (2018). Yleaf: Software for Human Y-Chromosomal Haplogroup Inference from Next-Generation Sequencing Data. Molecular Biology and Evolution, 35(5), 1291–1294.

Ralf, A., van Oven, M., Montiel González, D., de Knijff, P., van der Beek, K., Wootton, S., Lagacé, R., & Kayser, M. (2019). Forensic Y-SNP analysis beyond SNaPshot: High-resolution Y-chromosomal haplogrouping from low quality and quantity DNA using Ion AmpliSeq and targeted massively parallel sequencing. Forensic Science International. Genetics, 41, 93–106.

The Y Chromosome Consortium. (2002). A Nomenclature System for the Tree of Human Y-Chromosomal Binary Haplogroups. In Genome Research (Vol. 12, Issue 2, pp. 339–348). https://doi.org/10.1101/gr.217602

Underhill, P. A., & Kivisild, T. (2007). Use of y chromosome and mitochondrial DNA population structure in tracing human migrations. Annual Review of Genetics, 41, 539–564.

Van Geystelen, A., Decorte, R., & Larmuseau, M. H. D. (2013). AMY-tree: an algorithm to use whole genome SNP calling for Y chromosomal phylogenetic applications. BMC Genomics, 14, 101.

Zeng, T. C., Aw, A. J., & Feldman, M. W. (2018). Cultural hitchhiking and competition between patrilineal kin groups explain the post-Neolithic Y-chromosome bottleneck. Nature Communications, 9(1), 2077.

Zhang, F., Chen, R., Liu, D., Yao, X., Li, G., Jin, Y., Yu, C., Li, Y., & Coin, L. J. M. (2013). YHap: a population model for probabilistic assignment of Y haplogroups from re-sequencing data. BMC Bioinformatics, 14, 331.

Zhou, Z., Zhou, Y., Li, Z., Yao, Y., Yang, Q., Qian, J., Shao, C., Qian, X., Sun, K., Tang, Q., & Xie, J. (2022). Identification and assessment of a subset of Y-SNPs with recurrent mutation for forensic purpose. Forensic Science International, 334, 111270.

