## Supplementary Material - Tables for "A benchmarking of human Y-chromosomal haplogroup classifiers from whole-genome and whole-exome sequence data"

### Supplementary Materials

**Table S1.** Short-read and long-read sequencing summaries of the samples.

| | *Whole-Genome Sequencing Illumina* | | | | | *Whole-Exome Sequencing Illumina* | | | | | *Whole-Genome Sequencing Oxford Nanopore* | | | |
|---|---|---|---|---|---|---|---|---|---|---|---|---|---|---|
| ***Samples*** | ***Library Preparation*** | ***Reads*** | ***Mean Depth of Coverage*** | ***Mean Depth per SNV*** | ***% NRY covered*** | ***Library Preparation*** | ***Reads*** | ***Mean Depth of Coverage*** | ***Mean Depth per SNV*** | ***% NRY covered*** | ***Library Preparation*** | ***Reads*** | ***Mean Depth of Coverage*** | ***% NRY covered*** |
| **ATTA022** | Nextera-DNA-Prep | 6,201,454 | 10 | 47 | 28 | Illumina DNA Prep with Enrichment | 307,731 | 49 | 32 | 69 | - | - | - | - |
| **BLA-050** | Nextera-DNA-Prep | 8,890,179 | 14 | 73 | 39 | Illumina DNA Prep with Enrichment | 534,379 | 96 | 58 | 91 | - | - | - | - |
| **CAM-GFF-117** | Nextera-DNA-Prep | 4,429,452 | 7 | 36 | 19 | Illumina DNA Prep with Enrichment | 370,722 | 69 | 34 | 79 | - | - | - | - |
| **CAM-PYKL-042** | Nextera-DNA-Prep | 10,896,378 | 17 | 76 | 40 | Illumina DNA Prep with Enrichment | 528,802 | 95 | 46 | 83 | - | - | - | - |
| **CAM-PYKL-083** | Nextera-DNA-Prep | 6,244,694 | 10 | 49 | 29 | Illumina DNA Prep with Enrichment | 508,906 | 82 | 38 | 77 | - | - | - | - |
| **CAM-PYKL-102** | Nextera-DNA-Prep | 5,116,461 | 8 | 42 | 21 | Illumina DNA Prep with Enrichment | 439,601 | 69 | 36 | 72 | - | - | - | - |
| **CAM-PYKL-140** | Nextera-DNA-Prep | 10,009,468 | 15 | 70 | 39 | Illumina DNA Prep with Enrichment | 601,355 | 111 | 49 | 84 | - | - | - | - |
| **CAM-PYKL-196** | Nextera-DNA-Prep | 6,953,783 | 12 | 55 | 37 | Illumina DNA Prep with Enrichment | 508,129 | 88 | 42 | 86 | - | - | - | - |
| **CANIS0005** | Illumina-DNA-Prep | 7,733,041 | 10 | 29 | 28 | Nextera DNA Exome | 602,741 | 54 | 21 | 86 | - | - | - | - |
| **CANIS0018** | Illumina-DNA-Prep | 6,861,802 | 9 | 59 | 24 | Nextera DNA Exome | 684,540 | 74 | 27 | 93 | SQK LSK109 library prep | 243,552 | 14 | 39 |
| **CANIS0025** | Illumina-DNA-Prep | 18712,303 | 28 | 10 | 40 | Nextera DNA Exome | 561,182 | 60 | 26 | 87 | - | - | - | - |
| **CANIS0031** | Illumina-DNA-Prep | 8,225,012 | 13 | 71 | 38 | Nextera DNA Exome | 721,281 | 64 | 28 | 93 | - | - | - | - |
| **CANIS0057** | Nextera-DNA-Prep | 10,390,095 | 17 | 69 | 40 | Nextera DNA Exome | 535,888 | 57 | 32 | 80 | - | - | - | - |
| **CANIS0897** | Nextera-DNA-Prep | 7,598,324 | 13 | 47 | 33 | Nextera DNA Exome | 373,294 | 44 | 21 | 76 | - | - | - | - |
| **CANIS0123** | Illumina-DNA-Prep | 4,094,921 | 6 | 38 | 13 | Nextera DNA Exome | 161,617 | 27 | 20 | 62 | - | - | - | - |
| **CANIS0140** | Illumina-DNA-Prep | 6,398,874 | 6 | 34 | 12 | Nextera DNA Exome | 917,472 | 99 | 26 | 94 | SQK LSK109 library prep | 155,743 | 10 | 28 |
| **CANIS0141** | Illumina-DNA-Prep | 7,600,080 | 13 | 60 | 36 | Nextera DNA Exome | 659,253 | 66 | 24 | 92 | - | - | - | - |
| **CANIS0166** | Nextera-DNA-Prep | 9,139,067 | 15 | 63 | 38 | Nextera DNA Exome | 642,086 | 57 | 29 | 87 | - | - | - | - |
| **CANIS0170** | Illumina-DNA-Prep | 9,297,218 | 14 | 76 | 36 | Nextera DNA Exome | 679,582 | 62 | 27 | 91 | - | - | - | - |
| **CANIS0248** | Nextera-DNA-Prep | 8,168,328 | 14 | 58 | 39 | Nextera DNA Exome | 881,277 | 79 | 25 | 92 | - | - | - | - |
| **CANIS0259** | Illumina-DNA-Prep | 4,328,561 | 6 | 40 | 16 | Nextera DNA Exome | 566,683 | 57 | 19 | 85 | - | - | - | - |
| **CANIS0278** | Illumina-DNA-Prep | 8,497,448 | 13 | 67 | 35 | Nextera DNA Exome | 572,487 | 59 | 20 | 87 | - | - | - | - |
| **CANIS0295** | Illumina-DNA-Prep | 9,391,412 | 13 | 71 | 37 | Nextera DNA Exome | 790,743 | 79 | 27 | 93 | SQK LSK109 library prep | 209,400 | 12 | 36 |
| **CANIS0298** | Illumina-DNA-Prep | 10,903378 | 17 | 69 | 40 | Nextera DNA Exome | 819,128 | 77 | 22 | 87 | - | - | - | - |
| **CANIS0304** | Nextera-DNA-Prep | 8,786369 | 15 | 68 | 40 | Nextera DNA Exome | 594,938 | 58 | 23 | 90 | - | - | - | - |
| **CANIS0342** | Nextera-DNA-Prep | 10,767,293 | 17 | 72 | 40 | Nextera DNA Exome | 505,919 | 50 | 25 | 87 | - | - | - | - |
| **CANIS0360** | Illumina-DNA-Prep | 5,600,141 | 9 | 52 | 29 | Nextera DNA Exome | 708,873 | 76 | 29 | 92 | - | - | - | - |
| **CANIS0363** | Illumina-DNA-Prep | 8,222,187 | 14 | 63 | 37 | Nextera DNA Exome | 792,584 | 83 | 27 | 93 | SQK LSK109 library prep | 196,481 | 12 | 31 |
| **CANIS0369** | Nextera-DNA-Prep | 13,720,032 | 22 | 86 | 40 | Nextera DNA Exome | 691,589 | 64 | 24 | 82 | - | - | - | - |
| **CANIS0376** | Illumina-DNA-Prep | 8,703,404 | 12 | 74 | 37 | Nextera DNA Exome | 177,934 | 30 | 24 | 65 | - | - | - | - |
| **CANIS0419** | Nextera-DNA-Prep | 9,928,408 | 17 | 98 | 41 | Nextera DNA Exome | 531,336 | 55 | 20 | 87 | - | - | - | - |
| **CANIS0461** | Nextera-DNA-Prep | 8,322,400 | 14 | 67 | 39 | Nextera DNA Exome | 340,853 | 37 | 19 | 80 | - | - | - | - |
| **CANIS0501** | Illumina-DNA-Prep | 7,319,097 | 13 | 64 | 39 | Nextera DNA Exome | 694,615 | 75 | 30 | 92 | - | - | - | - |
| **CANIS0540** | Illumina-DNA-Prep | 7,959,917 | 14 | 66 | 38 | Nextera DNA Exome | 656,907 | 63 | 27 | 91 | - | - | - | - |
| **CANIS0567** | Nextera-DNA-Prep | 9,524,504 | 16 | 58 | 39 | Nextera DNA Exome | 543,182 | 45 | 19 | 85 | - | - | - | - |
| **CANIS0591** | Illumina-DNA-Prep | 8,819,247 | 12 | 41 | 35 | Nextera DNA Exome | 606,045 | 62 | 22 | 85 | - | - | - | - |
| **CANIS0630** | Nextera-DNA-Prep | 9,438,745 | 16 | 90 | 38 | Illumina DNA Prep with Enrichment | 563,814 | 83 | 51 | 84 | - | - | - | - |
| **CANIS0664** | Illumina-DNA-Prep | 11,057,346 | 15 | 78 | 37 | Nextera DNA Exome | 880,939 | 81 | 32 | 94 | SQK LSK109 library prep | 127,364 | 8 | 20 |
| **CANIS0686** | Illumina-DNA-Prep | 8,607,509 | 12 | 40 | 34 | Nextera DNA Exome | 571,960 | 58 | 39 | 75 | - | - | - | - |
| **CANIS0688** | Illumina-DNA-Prep | 9,795,141 | 12 | 53 | 24 | Nextera DNA Exome | 292,213 | 39 | 29 | 67 | - | - | - | - |
| **CANIS0700** | Nextera-DNA-Prep | 7,556,693 | 14 | 77 | 40 | Nextera DNA Exome | 519,874 | 48 | 21 | 85 | - | - | - | - |
| **CANIS0705** | Illumina-DNA-Prep | 7,241,520 | 11 | 68 | 34 | Nextera DNA Exome | 819,291 | 91 | 35 | 94 | SQK LSK109 library prep | 151,730 | 11 | 32 |
| **CANIS0708** | Illumina-DNA-Prep | 5,906,851 | 10 | 54 | 32 | Nextera DNA Exome | 597,745 | 58 | 18 | 84 | - | - | - | - |
| **CANIS0781** | Illumina-DNA-Prep | 7,952,665 | 9 | 64 | 30 | Nextera DNA Exome | 278,520 | 44 | 30 | 72 | SQK LSK109 library prep | 157,784 | 11 | 35 |
| **CANIS0864** | Nextera-DNA-Prep | 9,430,010 | 16 | 80 | 40 | Nextera DNA Exome | 619,172 | 60 | 26 | 91 | - | - | - | - |
| **GOU044** | Nextera-DNA-Prep | 8,287,525 | 14 | 65 | 38 | Illumina DNA Prep with Enrichment | 566,886 | 85 | 43 | 77 | - | - | - | - |
| **MD-22** | Nextera-DNA-Prep | 6,940,271 | 12 | 59 | 38 | Illumina DNA Prep with Enrichment | 348,789 | 66 | 35 | 84 | - | - | - | - |
| **NA24385** | Illumina-DNA-Prep | 8,696,533 | 15 | 85 | 40 | Illumina DNA Prep with Enrichment | 828,536 | 94 | 109 | 90 | - | - | - | - |
| **SOK9** | Nextera-DNA-Prep | 4,160,960 | 7 | 35 | 10 | Illumina DNA Prep with Enrichment | 439,254 | 72 | 45 | 83 | - | - | - | - |
| **WAZ118** | Nextera-DNA-Prep | 7,666,855 | 13 | 59 | 37 | Illumina DNA Prep with Enrichment | 489,425 | 92 | 48 | 88 | - | - | - | - |

**Table S2.** Per sample Y chromosome haplogroup classifications for the five tools based on short-read data. The retrieved classification that matches the consensus is shown in bold. A) WGS and B) WES datasets.

| Samples | Consensus haplogroup | Concordance rate (WGS) | Consensus haplogroup corrected | Whole-Genome Sequencing | | | | | | Whole-Exome Sequencing | | | | | |
|---|---|---|---|---|---|---|---|---|---|---|---|---|---|---|---|
| | | | | YLeaf (BAM) | Haplogrouper (BAM) | LineageTracker (BAM) | LineageTracker (VCF) | clean_tree_v2 (BAM) | pathPhynder (BAM) | YLeaf (BAM) | Haplogrouper (BAM) | LineageTracker (BAM) | LIneageTracker (VCF) | clean_tree_v2 (BAM) | pathPhynder (BAM) |
| ATTA022 | E1b1a1a1a2a1a3b1a19 | 100.00 | E1b1a1a1a2a1a3b1a19 | **E1b1a1a1a2a1a3b1a19** | **E1b1a1a1a2a1a3b1a19** | **E1b1a1a1a2a1a3b1a19** | **E1b1a1a1a2a1a3b1a19** | **E1b1a1a1a2a1a3b1a19** | **E1b1a1a1a2a1a3b1a19** | **E1b1a1a1a2a1a3b1a19** | **E1b1a1a1a2a1a3b1a19** | No Data | **E1b1a1a1a2a1a3b1a19** | **E1b1a1a1a2a1a3b1a19** | **E1b1a1a1a2a1a3b1a19** |
| BLA-050 | R1b1b2a2a1 | 100.00 | R1b1b2a2a1 | **R1b1b2a2a1** | **R1b1b2a2a1** | **R1b1b2a2a1** | **R1b1b2a2a1** | **R1b1b2a2a1** | **R1b1b2a2a1** | **R1b1b2a2a1** | R1 | No Data | P1orK2b2a | **R1b1b2a2a1** | **R1b1b2a2a1** |
| CAM-GFF-117 | E1b1a1a1a1c2 | 33.33 | E1b1a1a1a1c2 | **E1b1a1a1a1c2** | **E1b1a1a1a1c2**a | **E1b1a1a1a1c2**c3a1b | **E1b1a1a1a1c2**a | **E1b1a1a1a1c2**c3a1b | I1a2a1a1a2b3a | **E1b1a1a1a1c2** | E1b1a1a1 | No Data | E1b1a1a1 | **E1b1a1a1a1c2** | I1a2a1a1a2b3a |
| CAM-PYKL-042 | E1b1a1a1a1c1a1a3a1a1 | 100.00 | E1b1a1a1a1c1a1a3a1 | **E1b1a1a1a1c1a1a3a1**a1 | **E1b1a1a1a1c1a1a3a1**a1 | **E1b1a1a1a1c1a1a3a1**a1 | **E1b1a1a1a1c1a1a3a1**a1 | **E1b1a1a1a1c1a1a3a1**a1 | **E1b1a1a1a1c1a1a3a1**a1 | **E1b1a1a1a1c1a1a3a1** | E1b1a1a1a1c1a1 | No Data | E1b1a1a1a1c1a1 | **E1b1a1a1a1c1a1a3a1** | **E1b1a1a1a1c1a1a3a1** |
| CAM-PYKL-083 | E1b1a1a1a1c1a1a3l | 100.00 | E1b1a1a1a1c1a1a | **E1b1a1a1a1c1a1a**3l | **E1b1a1a1a1c1a1a**3l | **E1b1a1a1a1c1a1a**3l | **E1b1a1a1a1c1a1a**3l | **E1b1a1a1a1c1a1a**3l | **E1b1a1a1a1c1a1a**3l | **E1b1a1a1a1c1a1a** | E1b1a1a1a1c1a1 | No Data | E1b1a1a1a1c1a1 | **E1b1a1a1a1c1a1a** | **E1b1a1a1a1c1a1a** |
| CAM-PYKL-102 | E1b1a1a1a2a1a3a2b1 | 50.00 | E1b1a1a1a2a1a3a | **E1b1a1a1a2a1a3a**2b | **E1b1a1a1a2a1a3a**2 | **E1b1a1a1a2a1a3a**2b1 | **E1b1a1a1a2a1a3a**2 | **E1b1a1a1a2a1a3a**2b | **E1b1a1a1a2a1a3a**2b1 | **E1b1a1a1a2a1a3a** | E1b1a1a1 | No Data | E1b1a1a1 | **E1b1a1a1a2a1a3a** | **E1b1a1a1a2a1a3a** |
| CAM-PYKL-140 | E1b1a1a1a1c1a1a3l | 100.00 | E1b1a1a1a1c1a1a3l | **E1b1a1a1a1c1a1a3l** | **E1b1a1a1a1c1a1a3l** | **E1b1a1a1a1c1a1a3l** | **E1b1a1a1a1c1a1a3l** | **E1b1a1a1a1c1a1a3l** | **E1b1a1a1a1c1a1a3l** | E1b1a1a1a1c1a1a3a1d1b1b1a2 | E1b1a1a1 | No Data | E1b1a1a1 | E1b1a1a1a1c1a1a3a1d1b1b1a2 | **E1b1a1a1a1c1a1a3l** |
| CAM-PYKL-196 | E1b1a1a1a2a1a3b1d1c1a | 100.00 | E1b1a1a1a2a1a3b1d1c1a | **E1b1a1a1a2a1a3b1d1c1a** | **E1b1a1a1a2a1a3b1d1c1a** | **E1b1a1a1a2a1a3b1d1c1a** | **E1b1a1a1a2a1a3b1d1c1a** | **E1b1a1a1a2a1a3b1d1c1a** | **E1b1a1a1a2a1a3b1d1c1a** | **E1b1a1a1a2a1a3b1d1c1a** | E1b1a1a1 | No Data | E1b1a1a1 | **E1b1a1a1a2a1a3b1d1c1a** | **E1b1a1a1a2a1a3b1d1c1a** |
| CANIS0005 | J2a1 | 83.33 | J2a1 | **J2a1** | **J2a1** | **J2a1**a1b3 | **J2a1** | **J2a1** | **J2a1** | **J2a1** | J2a | No Data | J2 | **J2a1** | **J2a1** |
| CANIS0018 | R1b1a1b1a1a2a5 | 83.33 | R1b1a1b1a1a | **R1b1a1b1a1a**2a5 | **R1b1a1b1a1a**2a5 | **R1b1a1b1a1a**2c1a4b4a1a | **R1b1a1b1a1a**2a5 | **R1b1a1b1a1a**2a5 | **R1b1a1b1a1a**2a5 | **R1b1a1b1a1a** | R1 | No Data | P1orK2b2a | **R1b1a1b1a1a** | **R1b1a1b1a1a** |
| CANIS0025 | J1a2a1a2d2b2a1c | 33.33 | J1a2a1a2d2b2 | **J1a2a1a2d2b2**a1c | J1a2a1a2b | **J1a2a1a2d2b2**b2c4b1c3a | **J1a2a1a2d2b2**a1c1 | **J1a2a1a2d2b2**a1c1 | **J1a2a1a2d2b2** | **J1a2a1a2d2b2**b2c4d2a2a1c1 | J | No Data | J | **J1a2a1a2d2b2**b2c4d2a2a1c1 | **J1a2a1a2d2b2** |
| CANIS0031 | R1b1a1b1a1a2a | 66.67 | R1b1a1b1a1a2a | **R1b1a1b1a1a2a** | **R1b1a1b1a1a2a** | R1b1a1b1a1a2c1a2a2d1a | **R1b1a1b1a1a2a** | R1b1a1b1a1a | **R1b1a1b1a1a2a** | R1b1a1b1a1a2c1a4a1 | R1 | No Data | P1orK2b2a | R1b1a1b1a1a2c1a4a1 | R1b1a1b1a1a2c1a1a1a1a1 |
| CANIS0057 | R1b1a1b1a1a1c1a1 | 100.00 | R1b1a1b1a1a1c1a1 | **R1b1a1b1a1a1c1a1** | **R1b1a1b1a1a1c1a1** | **R1b1a1b1a1a1c1a1** | **R1b1a1b1a1a1c1a1** | **R1b1a1b1a1a1c1a1** | **R1b1a1b1a1a1c1a1** | R1b1a1b1a1a2b1c1a1a1 | R1 | No Data | P1orK2b2a | R1b1a1b1a1a2b1c1a1a1 | R1b1a1b1a1a2b1c1a1a1 |
| CANIS0897 | J2a1a1a2b2a2b2b1 | 66.67 | J2a1a1a2b2a2b | **J2a1a1a2b2a2b**2b1 | **J2a1a1a2b2a2b** | **J2a1a1a2b2a2b**2b1 | **J2a1a1a2b2a2b**2b1 | **J2a1a1a2b2a2b**2b1 | **J2a1a1a2b2a2b**2 | **J2a1a1a2b2a2b** | J2a1a | No Data | J2a1a | **J2a1a1a2b2a2b** | **J2a1a1a2b2a2b** |
| CANIS0123 | R1a1a1b2a2a3 | 33.33 | R1a1a1 | **R1a1a1**b2a2a3 | **R1a1a1**b2a2 | **R1a1a1**b2a1a2c1a | **R1a1a1**b2a2a3b | **R1a1a1**b2a2a3 | **R1a1a1**b2a2a3b | **R1a1a1** | R1 | No Data | I1a2a1a1a3a2 | **R1a1a1** | **R1a1a1** |
| CANIS0140 | G2a2b2a1a1c1a1a2a1a1 | 100.00 | G2a2b2a1a1c1a1 | **G2a2b2a1a1c1a1**a2a1a1 | **G2a2b2a1a1c1a1**a2a1a1 | **G2a2b2a1a1c1a1**a2a1a1 | **G2a2b2a1a1c1a1**a2a1a1 | **G2a2b2a1a1c1a1**a2a1a1 | **G2a2b2a1a1c1a1**a2a1a1 | **G2a2b2a1a1c1a1** | **G2a2b2a1a1c1a1** | No Data | G2a2b2a | **G2a2b2a1a1c1a1** | **G2a2b2a1a1c1a1** |
| CANIS0141 | J2b2a1a1a1a1a1a1c | 83.33 | J2b2a1a1a1a1a1a1c | **J2b2a1a1a1a1a1a1c** | J2b2a1a1a1a1a1a | **J2b2a1a1a1a1a1a1c** | **J2b2a1a1a1a1a1a1c** | **J2b2a1a1a1a1a1a1c** | **J2b2a1a1a1a1a1a1c** | **J2b2a1a1a1a1a1a1c** | J2 | No Data | J2 | **J2b2a1a1a1a1a1a1c** | J2b2a1a1a1a1a1a1 |
| CANIS0166 | R1a1a1b1a2b3a3a1a1 | 83.33 | R1a1a1b1a2b3a3a1a1 | **R1a1a1b1a2b3a3a1a1** | **R1a1a1b1a2b3a3a1a1** | **R1a1a1b1a2b3a3a1a1** | **R1a1a1b1a2b3a3a1a1** | **R1a1a1b1a2b3a3a1a1** | R1a1a1b1a2b3a3a1a | **R1a1a1b1a2b3a3a1a1** | R1 | No Data | I1a2a1a1a3a2 | **R1a1a1b1a2b3a3a1a1** | R1a1a1b1a2b3a3a1a |
| CANIS0170 | I1a3a1a1 | 66.67 | I1a3a1a1 | **I1a3a1a1** | **I1a3a1a1** | **I1a3a1a1**b | **I1a3a1a1**b | **I1a3a1a1**b | **I1a3a1a1** | **I1a3a1a1** | I1 | No Data | I | **I1a3a1a1** | I1 |
| CANIS0248 | T1a2a | 100.00 | T1a2a | **T1a2a** | **T1a2a** | **T1a2a** | **T1a2a** | **T1a2a** | **T1a2a** | **T1a2a** | T1a | No Data | TorK1b | **T1a2a** | **T1a2a** |
| CANIS0259 | E1b1b1a1a1c1a | 50.00 | E1b1b1a1a1c1a | **E1b1b1a1a1c1a** | **E1b1b1a1a1c1a** | E1b1b1a1a1c1b1a1a1 | E1b1b1a1a1c1b1a1a1 | E1b1b1a1a1c1b1d | **E1b1b1a1a1c1a** | **E1b1b1a1a1c1a** | E1b1b1 | No Data | E1b1b | E1b1b1a1a1c1b1a | E1b1b1a1a1c1b |
| CANIS0278 | J1a2a1a2d2b2b2c4d2a2a5a1e2b1 | 66.67 | J1a2a1a2d2b2b2c4d2a2a5a | **J1a2a1a2d2b2b2c4d2a2a5a**1e2b1 | J1a2a1a2b | **J1a2a1a2d2b2b2c4d2a2a5a**1e2b1 | **J1a2a1a2d2b2b2c4d2a2a5a**1e2b1 | **J1a2a1a2d2b2b2c4d2a2a5a**1e2b1 | **J1a2a1a2d2b2b2c4d2a2a5a**1 | **J1a2a1a2d2b2b2c4d2a2a5a** | J | No Data | J | **J1a2a1a2d2b2b2c4d2a2a5a** | J1a2a1a2d2b2b2c4d2a2a |
| CANIS0295 | R1b1a1b1a1a2a1a1a1 | 83.33 | R1b1a1b1a1a2a1a1a | **R1b1a1b1a1a2a1a1a**1 | **R1b1a1b1a1a2a1a1a**1 | R1b1a1b1a1a2c1a2a2d1a | **R1b1a1b1a1a2a1a1a**1 | **R1b1a1b1a1a2a1a1a**1 | **R1b1a1b1a1a2a1a1a**1 | **R1b1a1b1a1a2a1a1a** | R1 | No Data | P1orK2b2a | **R1b1a1b1a1a2a1a1a** | **R1b1a1b1a1a2a1a1a** |
| CANIS0298 | G2a2b1b1a1b1e | 50.00 | G2a2b1b1a1b1e | **G2a2b1b1a1b1e** | G2a2b1b1a1b1 | G2a2b1b1a1b1a1b | **G2a2b1b1a1b1e** | **G2a2b1b1a1b1e** | G2a2b1b1a3 | **G2a2b1b1a1b1e** | R1 | No Data | . | **G2a2b1b1a1b1e** | **G2a2b1b1a1b1e** |
| CANIS0304 | R1b1a1b1a1a2c1a1i3a | 83.33 | R1b1a1b1a1a2c1a1i3a | **R1b1a1b1a1a2c1a1i3a** | **R1b1a1b1a1a2c1a1i3a** | R1b1a1b1a1a2c1a2a2d1a | **R1b1a1b1a1a2c1a1i3a** | **R1b1a1b1a1a2c1a1i3a** | **R1b1a1b1a1a2c1a1i3a** | **R1b1a1b1a1a2c1a1i3a** | I2a1a | No Data | P1orK2b2a | **R1b1a1b1a1a2c1a1i3a** | **R1b1a1b1a1a2c1a1i3a** |
| CANIS0342 | I2a1a1a1a1a | 50.00 | I2a1a1a1a1 | **I2a1a1a1a1**a | **I2a1a1a1a1**a | **I2a1a1a1a1**a4c3a1 | **I2a1a1a1a1**a1c5 | **I2a1a1a1a1**a | **I2a1a1a1a1**a5 | **I2a1a1a1a1** | R1 | No Data | I2a1a1a1 | **I2a1a1a1a1** | **I2a1a1a1a1** |
| CANIS0360 | R1b1a1b1a1a2a | 33.33 | R1b1a1b1a1a | **R1b1a1b1a1a**2a | **R1b1a1b1a1a**2a | **R1b1a1b1a1a**2c1a2a2d1a | **R1b1a1b1a1a**2a7 | **R1b1a1b1a1a** | **R1b1a1b1a1a**2a7 | **R1b1a1b1a1a** | G2a2b2a | No Data | P1orK2b2a | **R1b1a1b1a1a** | **R1b1a1b1a1a**2a7 |
| CANIS0363 | G2a2b2a1a1f | 66.67 | G2a2b2a1a1f | **G2a2b2a1a1f** | **G2a2b2a1a1f** | G2a2b2a1a1c2b1b4 | **G2a2b2a1a1f** | **G2a2b2a1a1f** | G2a2b2a1 | **G2a2b2a1a1f** | - | No Data | G2a2b2a | **G2a2b2a1a1f** | **G2a2b2a1a1f** |
| CANIS0369 | E1b1b1b1a1c | 83.33 | E1b1b1b1a1 | **E1b1b1b1a1**c | **E1b1b1b1a1**c | **E1b1b1b1a1**b3a | **E1b1b1b1a1**c | **E1b1b1b1a1**c | **E1b1b1b1a1**c | **E1b1b1b1a1** | **E1b1b1b1a1** | No Data | E1b1b | **E1b1b1b1a1** | **E1b1b1b1a1** |
| CANIS0376 | R1b1a1b1a1a2a5 | 83.33 | R1b1a1b1a1a | **R1b1a1b1a1a**2a5 | **R1b1a1b1a1a**2a5 | **R1b1a1b1a1a**2c1a2a2d1a | **R1b1a1b1a1a**2a5 | **R1b1a1b1a1a**2a5 | **R1b1a1b1a1a**2a5 | **R1b1a1b1a1a** | R1 | No Data | P1orK2b2a | **R1b1a1b1a1a** | **R1b1a1b1a1a** |
| CANIS0419 | R1b1a1b1a1a2a1a1a1a | 83.33 | R1b1a1b1a1a2a1a1a | **R1b1a1b1a1a2a1a1a**1a | **R1b1a1b1a1a2a1a1a**1a | R1b1a1b1a1a2c1a2a2d1a | **R1b1a1b1a1a2a1a1a**1a | **R1b1a1b1a1a2a1a1a**1a | **R1b1a1b1a1a2a1a1a**1a | R1b1a1b1a1a2b1c2b1a | R1 | No Data | P1orK2b2a | R1b1a1b1a1a2b1c2b1a | **R1b1a1b1a1a2a1a1a** |
| CANIS0461 | R1b1a1b1a1a2c1a1i | 83.33 | R1b1a1b1a1a2c1a1i | **R1b1a1b1a1a2c1a1i** | **R1b1a1b1a1a2c1a1i** | R1b1a1b1a1a2c1a3a2a1b1a3 | **R1b1a1b1a1a2c1a1i** | **R1b1a1b1a1a2c1a1i** | **R1b1a1b1a1a2c1a1i** | R1b1a1b1a1a2c1a1a1a1a1a1a1a1a1a2 | R1 | No Data | P1orK2b2a | R1b1a1b1a1a2c1a1a1a1a1a1a1a1a1a2 | **R1b1a1b1a1a2c1a1i** |
| CANIS0501 | R1b1a1b1a1a1c1a1 | 100.00 | R1b1a1b1a1a | **R1b1a1b1a1a**1c1a1 | **R1b1a1b1a1a**1c1a1 | **R1b1a1b1a1a**1c1a1 | **R1b1a1b1a1a**1c1a1 | **R1b1a1b1a1a**1c1a1 | **R1b1a1b1a1a**1c1a1 | **R1b1a1b1a1a** | R1 | No Data | P1orK2b2a | **R1b1a1b1a1a** | **R1b1a1b1a1a** |
| CANIS0540 | R1b1a1b1a1a2c1a1e | 66.67 | R1b1a1b1a1a2c1a1e | **R1b1a1b1a1a2c1a1e** | **R1b1a1b1a1a2c1a1e** | R1b1a1b1a1a2c1a4b4a1a | **R1b1a1b1a1a2c1a1e** | R1b1a1b1a1a2c | **R1b1a1b1a1a2c1a1e** | **R1b1a1b1a1a2c1a1e** | R1 | No Data | P1orK2b2a | R1b1a1b1a1a1c2b2b1a1a1b1 | R1b1a1b1a1a |
| CANIS0567 | E1b1b1b1a1 | 83.33 | E1b1b1b1a1 | **E1b1b1b1a1** | **E1b1b1b1a1** | **E1b1b1b1a1**b3a | **E1b1b1b1a1** | **E1b1b1b1a1** | **E1b1b1b1a1** | **E1b1b1b1a1** | **E1b1b1b1a1** | No Data | E1b1b | E1b1b1b1a2 | **E1b1b1b1a1** |
| CANIS0591 | R1b1a1b1a1a2c1a3 | 83.33 | R1b1a1b1a1a2c | **R1b1a1b1a1a2c**1a3 | **R1b1a1b1a1a2c**1a3 | **R1b1a1b1a1a2c**1a5d | **R1b1a1b1a1a2c**1a3 | **R1b1a1b1a1a2c**1a3 | **R1b1a1b1a1a2c**1a3 | **R1b1a1b1a1a2c**1a4b8 | R1 | No Data | P1orK2b2a | **R1b1a1b1a1a2c**1a4b8 | **R1b1a1b1a1a2c** |
| CANIS0630 | R1a1a1b1a2b3a3a1a1 | 83.33 | R1a1a1b1a2b3a3a1a | **R1a1a1b1a2b3a3a1a**1 | **R1a1a1b1a2b3a3a1a**1 | **R1a1a1b1a2b3a3a1a**1 | **R1a1a1b1a2b3a3a1a**1 | **R1a1a1b1a2b3a3a1a**1 | **R1a1a1b1a2b3a3a1a**1 | **R1a1a1b1a2b3a3a1a** | R1 | No Data | I1a2a1a1a3a2 | **R1a1a1b1a2b3a3a1a** | **R1a1a1b1a2b3a3a1a** |
| CANIS0664 | E1b1b1b1a1c | 100.00 | E1b1b1b1a1 | **E1b1b1b1a1**c | **E1b1b1b1a1**c | **E1b1b1b1a1**c | **E1b1b1b1a1**c | **E1b1b1b1a1**c | **E1b1b1b1a1**c | **E1b1b1b1a1** | **E1b1b1b1a1** | No Data | E1b1b | **E1b1b1b1a1** | **E1b1b1b1a1** |
| CANIS0686 | R1b1a1b1a1a2a5 | 83.33 | R1b1a1b1a1a2 | **R1b1a1b1a1a2**a5 | **R1b1a1b1a1a2**a5 | **R1b1a1b1a1a2**c1a2a2d1a | **R1b1a1b1a1a2**a5 | **R1b1a1b1a1a2**a5 | **R1b1a1b1a1a2**a5 | **R1b1a1b1a1a2** | R1 | No Data | P1orK2b2a | **R1b1a1b1a1a2** | **R1b1a1b1a1a2** |
| CANIS0688 | R1b1a1b1a1a2a | 66.67 | R1b1a1b1a1a | **R1b1a1b1a1a**2a | **R1b1a1b1a1a**2a | **R1b1a1b1a1a**2c1a1f1a1b | **R1b1a1b1a1a**2a | **R1b1a1b1a1a** | **R1b1a1b1a1a**2a | **R1b1a1b1a1a** | R1 | No Data | P1orK2b2a | **R1b1a1b1a1a**2c1a3a2a1b1 | **R1b1a1b1a1a** |
| CANIS0700 | R1b1a1b1a1a2a5 | 83.33 | R1b1a1b1a1a2 | **R1b1a1b1a1a2**a5 | **R1b1a1b1a1a2**a5 | **R1b1a1b1a1a2**c1a2a2d1a | **R1b1a1b1a1a2**a5 | **R1b1a1b1a1a2**a5 | **R1b1a1b1a1a2**a5 | **R1b1a1b1a1a2** | | No Data | P1orK2b2a | **R1b1a1b1a1a2**c1a2a2a1d1a | **R1b1a1b1a1a2** |
| CANIS0705 | R1b1a1b1a1a2b1a1 | 33.33 | R1b1a1b1a1a2 | **R1b1a1b1a1a2**b1a1 | **R1b1a1b1a1a2**b | **R1b1a1b1a1a2**c1a2a2d1a | **R1b1a1b1a1a2**b | **R1b1a1b1a1a2**b1a | **R1b1a1b1a1a2**b1a1 | **R1b1a1b1a1a2** | R1 | No Data | P1orK2b2a | R1b1a1b1a1a1c2b1a1a1 | **R1b1a1b1a1a2** |
| CANIS0708 | R1b1a1b1a1a2a1b1a1 | 50.00 | R1b1a1b1a1a2a1b1a1 | **R1b1a1b1a1a2a1b1a1** | **R1b1a1b1a1a2a1b1a1** | R1b1a1b1a1a2c1a2a2d1a | **R1b1a1b1a1a2a1b1a1** | **R1b1a1b1a1a2a1b1a1** | **R1b1a1b1a1a2a1b1a1**a | **R1b1a1b1a1a2a1b1a1** | R1 | No Data | P1orK2b2a | **R1b1a1b1a1a2a1b1a1** | E1b1a1a1a2a1a3b1d1c1b |
| CANIS0781 | T1a1a1b2b2b1a1a | 100.00 | T1a1a1b2b2b | **T1a1a1b2b2b**1a1a | **T1a1a1b2b2b**1a1a | **T1a1a1b2b2b**1a1a | **T1a1a1b2b2b**1a1a | **T1a1a1b2b2b**1a1a | **T1a1a1b2b2b**1a1a | **T1a1a1b2b2b** | **T1a1a1b2b2b** | No Data | TorK1b | **T1a1a1b2b2b** | **T1a1a1b2b2b** |
| CANIS0864 | R1b1a1b1a1a2a1a1a1 | 83.33 | R1b1a1b1a1a2a1a1a1 | **R1b1a1b1a1a2a1a1a1** | **R1b1a1b1a1a2a1a1a1** | R1b1a1b1a1a2c1a2a2d1a | **R1b1a1b1a1a2a1a1a1** | **R1b1a1b1a1a2a1a1a1** | **R1b1a1b1a1a2a1a1a1** | **R1b1a1b1a1a2a1a1a1** | R1 | No Data | P1orK2b2a | **R1b1a1b1a1a2a1a1a1** | **R1b1a1b1a1a2a1a1a1** |
| GOU044 | E1b1a1a1a2a1a3b1a | 100.00 | E1b1a1a1a2a1a3b1a | **E1b1a1a1a2a1a3b1a** | **E1b1a1a1a2a1a3b1a** | **E1b1a1a1a2a1a3b1a** | **E1b1a1a1a2a1a3b1a** | **E1b1a1a1a2a1a3b1a** | **E1b1a1a1a2a1a3b1a** | **E1b1a1a1a2a1a3b1a** | E1b1a1a1 | No Data | E1b1a1a1 | **E1b1a1a1a2a1a3b1a** | **E1b1a1a1a2a1a3b1a** |
| MD-22 | E1a1a2b | 100.00 | E1a1 | **E1a1**a2b | **E1a1**a2b | **E1a1**a2b | **E1a1**a2b | **E1a1**a2b | **E1a1**a2b | **E1a1** | E | No Data | E | **E1a1** | **E1a1** |
| NA24385 | J1a2a1a2c1a1a1 | 50.00 | J1a2a1a2c1a1a1 | J1a2a1a2c1a1a | J1a2a1a2c1a1 | **J1a2a1a2c1a1a1** | **J1a2a1a2c1a1a1** | J1a2a1a2c1a1a | **J1a2a1a2c1a1a1** | **J1a2a1a2c1a1a1** | J1a2a1a2c1a1 | No Data | J1a2a1a2c1a1 | **J1a2a1a2c1a1a1** | **J1a2a1a2c1a1a1** |
| SOK9 | R1b1b2a2b | 66.67 | R1b1b2a2b | R1b1b2a2 | **R1b1b2a2b** | **R1b1b2a2b** | **R1b1b2a2b** | R1b1b2a2 | **R1b1b2a2b** | R1b1b2a2 | R1 | No Data | P1orK2b2a | R1b1b2a2 | **R1b1b2a2b** |
| WAZ118 | E1b1a1a1a1c1b2a1 | 50.00 | E1b1a1a1a1c1b2a | **E1b1a1a1a1c1b2a**1 | **E1b1a1a1a1c1b2a**1 | **E1b1a1a1a1c1b2a**1b | **E1b1a1a1a1c1b2a**1b | **E1b1a1a1a1c1b2a**1 | **E1b1a1a1a1c1b2a**1b | **E1b1a1a1a1c1b2a** | E1b1a1a1 | No Data | E1b1a1a1 | **E1b1a1a1a1c1b2a** | **E1b1a1a1a1c1b2a** |
